# The topology of transmembrane protein-protein interaction interfaces is encoded in their physicochemical features

**DOI:** 10.64898/2026.07.16.738731

**Authors:** Lisa Allmesberger-Riegler, Fabian Frommelt, Brianda L. Santini, Giulio Superti-Furga, Evandro Ferrada, Peter Sykacek

## Abstract

Transmembrane (TM) protein-protein interactions (PPIs) are essential mediators of signal transduction, transport of solutes and communication, yet the biophysical features that characterize their diverse topologies remain poorly understood. To reduce this knowledge gap we use the human solute carrier (SLC) interactome to study whether physicochemical features of PPI interfaces encode information about their TM topology (i.e., their position and arrangement with respect to the membrane). To this end we predicted structures for 2, 055 experimentally validated PPIs using AlphaFold v3.0, performed molecular dynamics simulations and annotated the PPI interfaces by TM coverage. As a result every interface is characterized by 63 physicochemical, structural and energy features. A reproducible machine learning workflow allows us to study the interdependence between these interface properties and interface topology. We found that amino acid composition and secondary structure contributed most to distinguishing soluble from fully membrane-embedded interfaces. Membrane-embedded interfaces contained a larger fraction of hydrophobic residues and *α*-helices, while charged residues were depleted. For partially and full membrane-embedded interfaces, amino acid composition and secondary structure get more similar and differences are increasingly observed in charge and energy related features. As particularly noteworthy we find that PPI interface characteristics vary with the number of annotated TM segments mainly through differences in proportions of secondary structure, charge, and flexibility. We hence conclude that PPI interface characteristics harbor substantial information about TM interface topology and provide a framework for the study and design of membrane protein interaction interfaces.

## 1 INTRODUCTION

Protein-protein interactions (PPIs) are fundamental to cellular physiology, mediating processes that range from signal transduction to metabolic regulation (Stumpf et al. 2008, Rual et al. 2005). Transmembrane (TM) PPIs are particularly relevant, because they mediate communication across lipid bilayers, sense and amplify cellular signals, and transport metabolites (Schaller and Lauschke (2019), Superti-Furga et al. (2020)).

Dedicated interactomics and proximity labeling studies have mapped extensive interaction networks for several major classes of membrane-bound proteins, including receptor tyrosine kinases (RTKs), G-protein-coupled receptors (GPCRs), and solute carriers (SLCs) (Salokas et al. 2022, Polacco et al. 2024, Kotliar et al. 2024, Sokolina et al. 2017, Frommelt et al. 2025).

Despite these experimental efforts and their biological relevance, TM PPI interfaces remain substantially understudied. A recent estimate of the reference human interactome comprises approximately 117, 900 non-redundant PPIs, yet experimentally determined complex structures are available for only 3.95% of these interactions, with membrane proteins particularly underrepresented in structural datasets (Mosca et al. 2013, Kosoglu et al. 2024). However, the study of PPIs is shifting due to recent technological advancements in both experimental and computational methods.

Recent advancements in mass spectrometry (MS) data acquisition techniques for interactome mapping, together with the rise of AI, particularly AlphaFold (AF) (Jumper et al. 2021), have led to a large expansion of available structural models (Frommelt et al. 2026, Han et al. 2026). These computational methods have drastically increased the structural coverage of the human interactome, provided the opportunity to explore structure models of TM PPI interfaces and allowed the systematic study of TM PPIs and their interfaces (Abramson et al. 2024).

TM segments are known to have characteristic patterns of amino acid and secondary structure composition facing the lipid bilayer, and these patterns have been traditionally used to accurately predict TM proteins from sequence alone (von Heijne 1992, Cymer et al. 2015, Hallgren et al. 2022). However, it is unknown whether the same simple principles apply to sequence, structure or physicochemical determinants of TM PPI interfaces.

It is clear that depending on their arrangement with respect to the membrane, a PPI interface can have different topologies. For instance, they may be soluble or fully embedded within the membrane (Duarte et al. 2013). However, to the best of our knowledge, whether these topological differences between PPI interfaces can be predicted from physicochemical properties of the interface residues, has not been assessed. Addressing this problem requires resolving a variety of physicochemical features in an unbiased set of high-quality structure models of PPIs at atomic resolution.

Here, we explore this problem using the human solute carrier (SLC) interactome (Frommelt et al. 2025). This dataset contains PPIs involving SLCs, one of the largest superfamilies of TM transport proteins (Höglund et al. 2011). SLC transporters interact with both soluble and membrane proteins, making them well suited for studying a representative range of the topologies of PPI interfaces. We model 2, 055 experimentally supported PPIs, refine them by molecular dynamics simulations, and quantify the fraction of the interface coverage in the membrane or TM segment coverage per PPI interface.

We then use a suite of machine learning methods to investigate which physicochemical features are able to distinguish interfaces across different levels of membrane coverage and TM architectures. This provides us with a systematic framework to study the interplay between membrane topology and interface features, including energy related and geometric properties.

## 2 RESULTS

### 2.1 The topologies of TM PPI interfaces

TM PPI interfaces can adopt a variety of interaction interfaces. These range from fully soluble to deeply membrane-embedded PPI interfaces, with many cases falling somewhere in between (Duarte et al. 2013). Therefore, TM PPIs can be broadly grouped according to their topology into interactions that take place outside the membrane or fully embedded in the lipid bilayer (Figure 1a, b).

**Figure 1.**
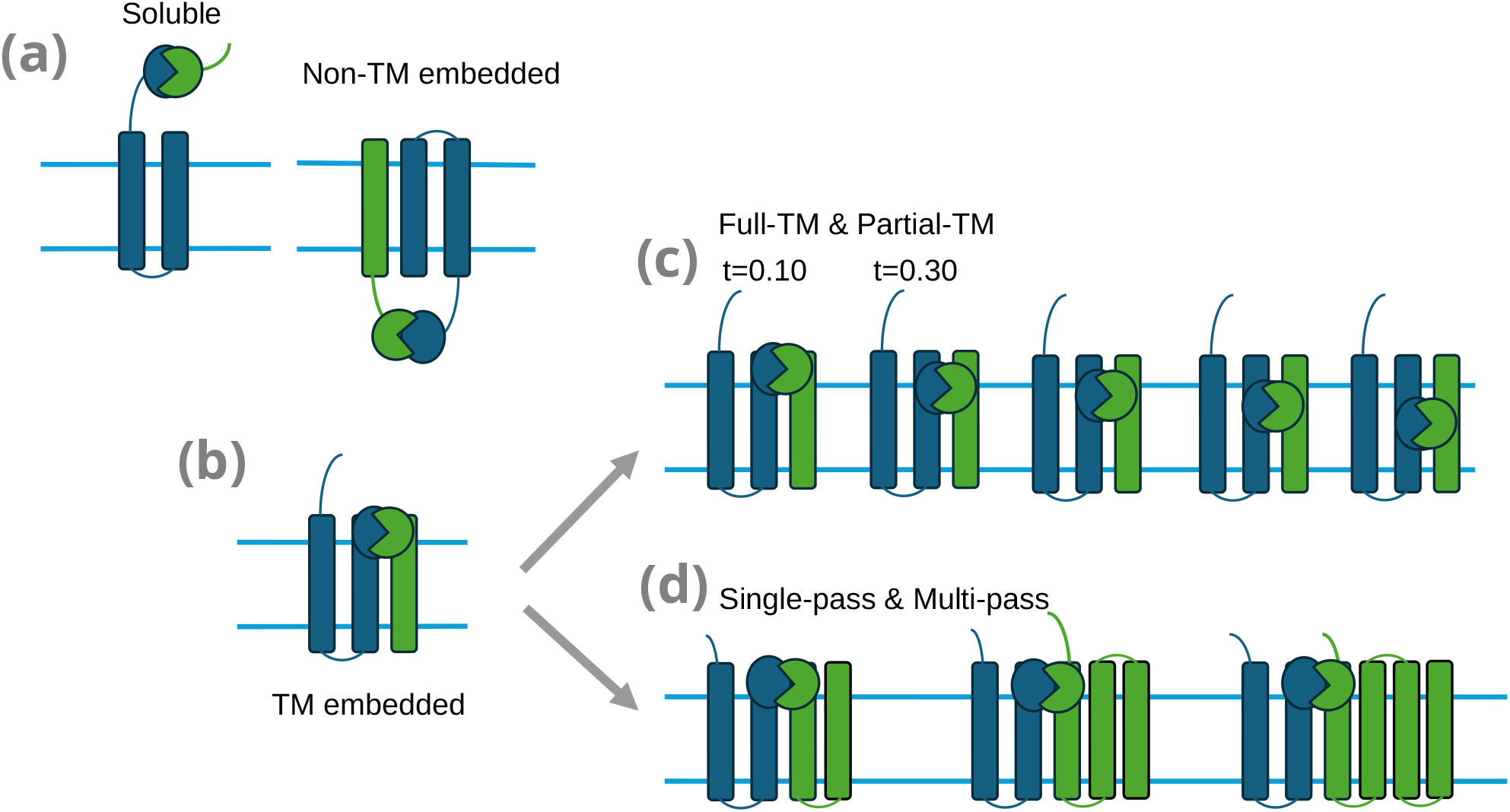
Schematic representation of TM PPI interface topologies. Blue and green helices represent the SLC transporter and its respective interactor. Horizontal lines indicate the membrane boundaries. a) A soluble interactor binding to the SLC outside of the membrane, representing cases with either no reported TM annotation or no interaction occurring with the TM domain. b) An interactor containing a TM domain interacting with the SLC through a soluble domain. c) Membrane-embedded interfaces of the interactor and the SLC with varying degrees of TM coverage. (D) Membrane-embedded interfaces involving interactors with single- or multi-pass TM domains.

PPI interfaces involving proteins with a TM domain might include interactions where one of the partners is not a membrane protein. These types of PPI interactions can occur at either side of the membrane, and engage either the N- or C-terminus of the TM protein (Figure 1a). In contrast, the topology of PPI interactions embedded in the membrane can show substantial heterogeneity. The interfaces of these PPIs might be partially or fully embedded in the membrane (Figure 1c) and involve one or multiple TM domains (Figure 1d). Whether TM PPI interfaces with different topologies are distinguishable by their physicochemical, structural, or energy features remains unknown.

### 2.2 Understanding the topologies of TM PPI interfaces

The goal of this work is to systematically characterize the heterogeneity of TM PPI interface topologies by exploring physicochemical, structural, and energy related features extracted from structural models of PPI interfaces. To achieve this, we utilized a dataset of nearly 19,000 experimentally validated binary PPIs, containing an SLC protein and a binding partner (interactor) (Frommelt et al. 2025). Structural prediction and confidence filtering yielded 2,055 high-confidence complexes for downstream analysis.

Although the SLC protein always contains at least one TM domain, the interactor is not necessarily a TM protein as described above (Figure 1a). Among the 2,055 PPIs in our dataset, approximately half (n=1,031) SLCs form a PPI with an interactor which is also a TM protein. The other half are soluble interactors (Figure 1a).

To fully characterize TM PPI topologies in our dataset, we retrieved TM domain annotations from UniProtKB (UniProt Consortium 2023) (Methods 4.3) and mapped these segments onto the residues of PPI interface. This allowed us to calculate the fraction of a PPI interface embedded in the membrane, or TM coverage, which we represent as the parameter *t*. This single parameter helped us distinguish between interface topologies with increasing degrees of membrane embedding. Under this metric, an interface with *t* = 0 is characterized as fully soluble, whereas *t* = 1.0 indicates an interface fully embedded in the membrane (Figure 1c).

Similarly, we used TM annotations from UniProtKB (UniProt Consortium 2023) to determine the number of TM segments in the interactor (Figure 1d). Interactors with exactly one TM segment are defined in this work as single-pass, whereas those with more than one TM segment are defined as multi-pass (Figure 1d).

To extract interface features, we generated structural models for all 19,000 experimentally obtained PPIs using AlphaFold v3.0 (AF3) (Abramson et al. 2024). AF3 models were filtered using confidence thresholds of ipTM *≥* 0.3 and pDockQ *≥* 0.23 (Burke et al. 2023), and each structure was refined by a 10 ns molecular dynamics (MD) simulation to stabilize the interface geometry (Methods 4.1). PPI interfaces were defined for the 2,055 filtered complexes based on a 10 Å distance cutoff. We used GetContacts (Venkatakrishnan et al. 2019) to extract contact residues, defining them as any residue present in more than 30% of the MD frames (30 out of 100 MD frames). Next, we extracted interface features per residue from the trimmed structures. These residue features were then aggregated over each PPI interface, resulting in 63 features per interface (Methods 4.2).

The 63 interface features span six broad categories that describe complementary physicochemical, structural, and energy related properties of PPI interfaces (Methods 4.2, Supplementary Table S1).

Here, Amino acid composition features describe the chemical characteristics of the interface through amino acid proportions. Secondary structure features describe the structural composition of the interface, distinguishing *α*-helical, *β*-sheet, and unordered (coil) structural content. Energy related terms derived from MM-GBSA calculations on MD trajectories, quantify binding energy contributions from MD trajectories. Geometric properties quantify the spatial compactness and planarity of the interface. Interface flexibility was quantified based on the root-mean squared fluctuation (RMSF) of residues. Finally, contact features describe the frequency and composition of intermolecular interaction types at the interface, such as salt bridges, hydrogen bonds, and aromatic interactions. These features were measured for each residue and contact in each of the PPI interfaces of our dataset. To prepare the data for further analyses, the features were aggregated, transformed and standardized (Methods 4.2).

In order to understand whether the features described above can be used to decode the diverse topologies of the TM PPI interface, we use a machine learning (ML) approach. We reason that different ML strategies can help us to determine whether combinations of these features can indeed distinguish between interface topologies, and by doing so, help us understand their biophysical properties. Specifically, we apply Support Vector Classifiers (SVC) (Vapnik (1998)), Multilayer Perceptrons (MLP) (Hornik (1991)), Random Forests (RF) classification (Breiman (2001)), and Gaussian Process Classifiers (GPC) (Rasmussen and Williams (2006)) and assess their performance employing accuracy (Acc) and area under the ROC curve (AUC) (Methods 4.5, 4.6). To identify interface features most distinct to specific topologies, we use feature importance analysis (Methods 4.7). The overall workflow is shown in Figure 2a. Figure 2b shows how we partition the dataset into topological classes, with representative complexes illustrating each case.

**Figure 2.**
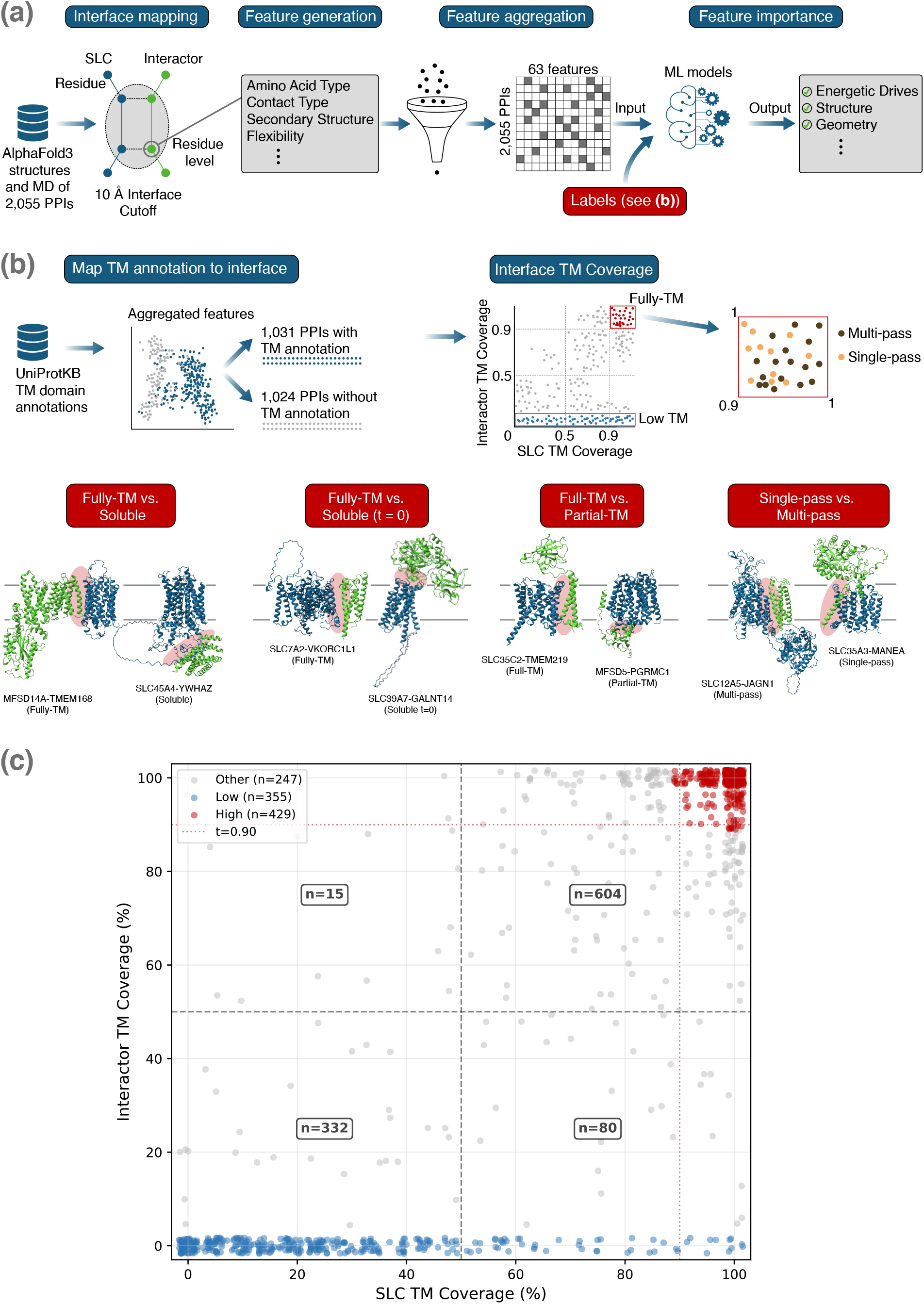
Overview of the analysis workflow, topological classification tasks, and TM-coverage distribution of interface residues. a) Schematic representation of the main data handling tasks used within the analysis workflow. High-confidence AlphaFold3 structures undergo iterative terminal trimming followed by 10 ns MD refinement to determine stability of interface residues. Residue features are extracted using a 10 Å heavy-atom distance cutoff and aggregated into a standardized 63-feature matrix across six biophysical categories. Combined with the Labels of Figure 2b, this matrix serves as the input for machine learning classification and feature importance analysis. b) Generation of interface labels. UniProtKB TM domain annotations are mapped onto interface residues, and interactions are grouped based on whether they are soluble or overlap with TM segments. This results in TM coverage values per interface. The high TM coverage group (>=0.90) is further annotated by the TM segments, allowing classification into single- and multi-pass categories. Structural models illustrate representative complexes for each topology, with the solute carrier (SLC) protein colored in blue and the interactor in green. The red shaded region outlines approximately the interaction interface. All protein coordinates represent trimmed structures and were rendered using UCSF ChimeraX (Meng et al. 2023). (c) Distribution of the interface TM coverage (%) for SLCs against their corresponding interactors (*n* = 1,031 pairs with TM annotation). Points are slightly jittered to prevent overlapping and enhance readability. Quadrant counts denote the number n of complexes falling within specified coverage boundaries. Red highlights the fully membrane-embedded PPIs. Blue dots are PPIs where the interactor contains zero TM coverage.

### 2.3 Amino acid composition and secondary structure robustly distinguish membrane-embedded from soluble interfaces

We begin by studying the two most general topologies of TM PPI interfaces, which are those that occur outside the membrane (i.e., soluble) or that are fully embedded in it (Figure 1a,b). Out of the 2, 055 PPI interfaces, 1, 031 had an interactor with an TM domain (annoated in UniProtKB) that overlaps with the interface residues. The remaining PPI interactors lack TM domain annotation in UniProtKB (Figure 2b. An overview of the TM coverage distribution is shown in Figure 2c, where we can see that many PPIs have high membrane coverage. We define fully embedded interfaces as those with *t ≥* 0.9 for both, SLC and interactor (Figure 1a). This group of interfaces contains 429 PPIs and is shown as red dots in Figure 2b,c. In contrast, the soluble group consists of interactors lacking TM domains.

To determine if soluble and fully embedded interfaces exhibit distinct characteristics in their features, we analyzed the 63 interface features based on four ML classifiers (SVC, MLP, RF, and GPC; Methods 4.5). These classifiers use different but complementary ways of separating topologies in the data. Every supervised learning method has its own bias (Wolpert 2002). So, using different classifiers allows us to assess whether the information captured by these classifiers reflects a consistent signal based on interface features (Methods 4.5).

All classifiers achieve 93–95% Acc and AUC values of up to 0.979 (Figure 3a, Table 1). This result indicates a strong physicochemical difference of interface features between soluble and fully embedded interfaces. Having established that the two topologies can be distinguished based on their interface features, we next ask which of these features are most informative. To answer this question, we use feature importance analysis which shows that amino acid composition and secondary structure are the most informative features, consistent with the physicochemical environment of the lipid bilayer (Figure 3b). In particular, the proportion of hydrophobic, positively and negatively charged residues ranked highest, followed by the proportion of coil and *α*-helix as well as the proportion of individual residue features (i.e., glutamic acid, aspartic acid, arginine and lysine).

**TABLE 1.** Classification performance for all four analyses and classifiers. Comparison of the different topologies, Clf: the used ML method; n indicates the number of total samples of both topologies combined; Acc: accuracy (%); AUC: area under ROC curve; MI: mutual information. Significance against majority-class baseline (McNemar’s test): ^*\*\*\**^*p* < 0.001; ^*\*\**^*p* < 0.01; ^*\**^*p* < 0.05; ns: not significant. RFC values reported as mean*±*std across 100 random subsampling runs.

| Comparison | Clf | n | Acc | AUC | MI | p |
| --- | --- | --- | --- | --- | --- | --- |
| Fully-TM ($t \geq 0.90$) vs. Soluble | SVC | 858 | 94.5 | 0.979 | 0.740 | *** |
|  | MLPC |  | 93.8 | 0.972 | 0.800 | *** |
|  | GPC |  | 93.8 | 0.977 | 0.396 | *** |
| | RFC | | 93.1±0.6 | 0.976±0.004 | 0.4037 | *** |
| Fully-TM ($t \geq 0.90$) vs. Soluble ($t=0$) | SVC | 710 | 94.9 | 0.991 | 0.780 | *** |
|  | MLPC |  | 95.4 | 0.988 | 0.851 | *** |
|  | GPC |  | 95.6 | 0.987 | 0.401 | *** |
| | RFC | | 95.3±0.4 | 0.987±0.001 | 0.550 | *** |
| full-TM vs. partial-TM ($t = 0.10$) | SVC | 492 | 90.7 | 0.972 | 0.631 | *** |
|  | MLPC |  | 91.9 | 0.953 | 0.703 | *** |
|  | GPC |  | 90.7 | 0.968 | 0.301 | *** |
| | RFC | | 90.3±0.8 | 0.965±0.004 | 0.404 | *** |
| full-TM vs. partial-TM ($t = 0.30$) | SVC | 292 | 84.3 | 0.922 | 0.397 | *** |
|  | MLPC |  | 83.2 | 0.900 | 0.428 | *** |
|  | GPC |  | 83.6 | 0.913 | 0.118 | *** |
| | RFC | | 83.6±1.6 | 0.918±0.009 | 0.214 | *** |
| full-TM vs. partial-TM ($t = 0.50$) | SVC | 190 | 76.8 | 0.830 | 0.190 | *** |
|  | MLPC |  | 73.7 | 0.783 | 0.290 | *** |
|  | GPC |  | 76.8 | 0.842 | 0.047 | *** |
| | RFC | | 76.1±3.2 | 0.841±0.025 | 0.124 | *** |
| full-TM vs. partial-TM ($t = 0.70$) | SVC | 204 | 66.3 | 0.75 | 0.087 | ** |
|  | MLPC |  | 65.8 | 0.70 | 0.200 | ** |
|  | GPC |  | 66.7 | 0.75 | 0.021 | ** |
| | RFC | | 68.3±3.3 | 0.755±0.032 | 0.044 | ** |
| full-TM vs. partial-TM ($t = 0.90$) | SVC | 314 | 55.4 | 0.560 | 0.003 | ns |
|  | MLPC |  | 59.8 | 0.620 | 0.150 | * |
|  | GPC |  | 59.8 | 0.630 | 0.004 | * |
| | RFC | | 60.2±2.1 | 0.640±0.027 | 0.014 | ** |
| Single-pass vs. Multi-pass | SVC | 429 | 82.7 | 0.895 | 0.342 | *** |
|  | MLPC |  | 81.4 | 0.863 | 0.505 | *** |
|  | GPC |  | 82.8 | 0.901 | 0.149 | *** |
|  | RFC |  | 82.1 | 0.894 | 0.205 | *** |

**Figure 3.**
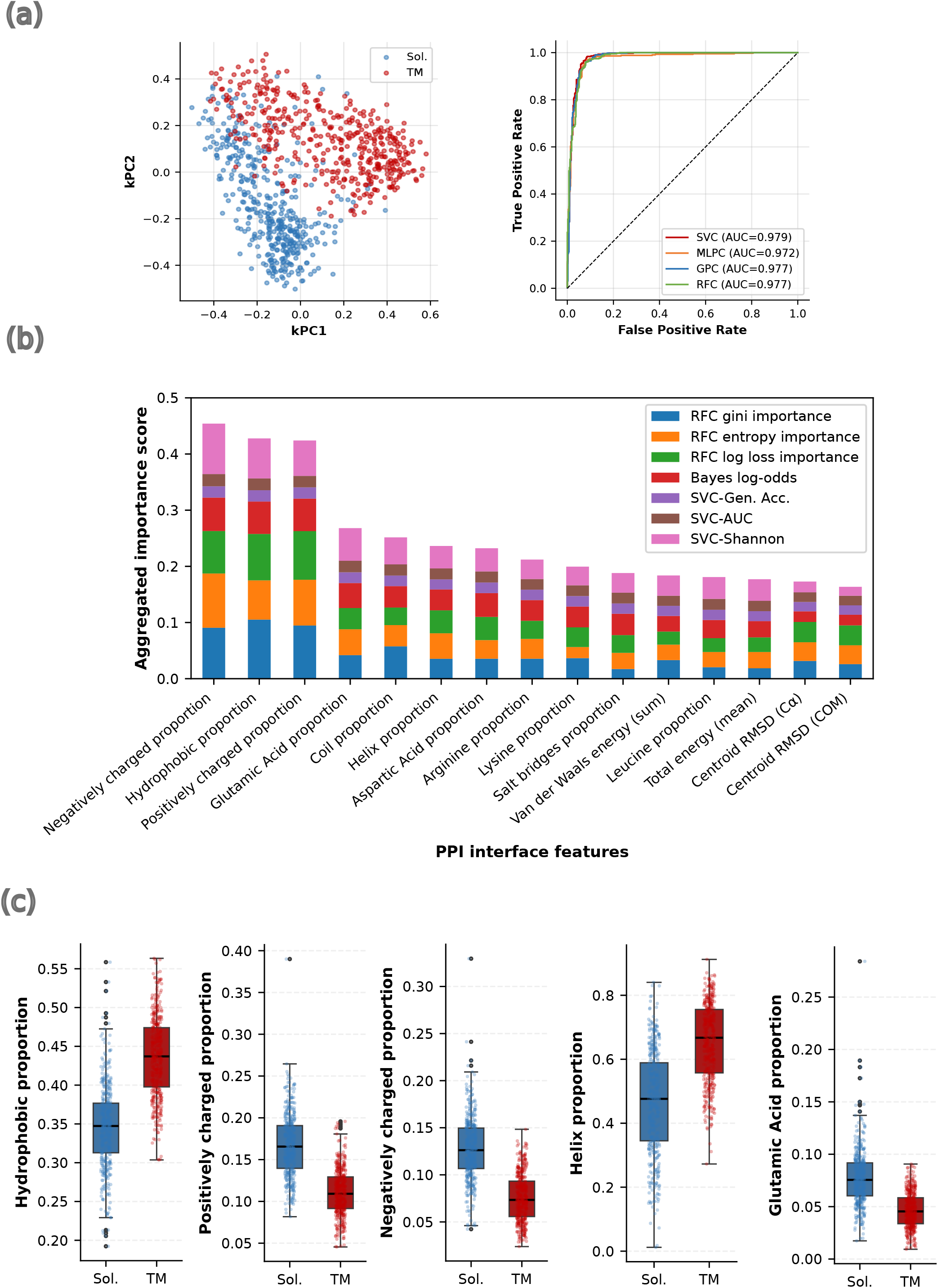
Physicochemical interface features distinguish membrane-embedded (TM) from soluble (Sol.) interfaces. a) Kernel PCA of interface feature space (left) and ROC curves with corresponding AUC values for four classifiers (right). (b) Aggregated feature importance scores for the top 15 interface features across seven importance metrics. c) Boxplots of the four top-ranked importance features. Blue: soluble interfaces (Sol.); Red: membrane-embedded interfaces (TM). Lower and upper hinges of box plots correspond to the 25th and 75th percentiles, respectively. Lower and upper whiskers extend from the hinge to the smallest or largest value no further than the 1.5× interquartile range from the hinge, respectively. Black line represents the median and the black dots represent outliers. All features shown are significant after FDR correction (adjusted p < 0.05).

Here, the proportion of hydrophobic residues is higher in membrane-embedded interfaces, reflecting the energy related favorability of non-polar side chains in the membrane core (Walters and DeGrado 2006).Proportions of positively and negatively charged residues are depleted at membrane-embedded interfaces (Figure 3c, Figure S1), consistent with the unfavorable cost of burying ionizable groups in the hydrophobic membrane environment (Cymer et al. 2015, von Heijne 1992). This trend is also reflected at the level of individual residues, with proportions of glutamic acid, aspartic acid, arginine, and lysine among the most strongly depleted features (Figure S1). Membrane-embedded interfaces also show lower proportion of coil and higher proportion of *α*-helices at the interface, consistent with the structural constraints of the lipid bilayer, which generally favor TM *α*-helices (von Heijne 1992).

To ensure that our conclusions are not affected by the absence of a TM domain in the interactors, we repeated the analysis using another sample of PPI interfaces. In this case, the interactors do have a TM domain but not at the interface (*t* = 0). The other topology is the same fully embedded dataset as before (*t* >= 0.90). Example PPI structures for this topology are shown in Figure 2b.

Classification performance in these datasets remained high, with 94–96% Acc and AUC values up to 0.991 (Table 1). This shows that the PPI interface features can distinguish between these topologies and differ among them. The same major features remained the most informative, including proportions of hydrophobicity, charge, and secondary structure, although their relative contributions shifted modestly. In particular, structural features became slightly more important, while the proportion of charged residues such as glutamic acid contributed less strongly (Figure S2, S3). This may indicate that the proportions of charged residues at the interface partly reflect broader differences between soluble and TM protein environments

Together, these results show that soluble versus fully embedded PPI interface topologies leave a clear physicochemical signal in the composition of PPI interfaces. And this signal is driven by the strong energetic constraint of the lipid bilayer versus the soluble environment.

### 2.4 Physicochemical distinctions weaken with increasing TM coverage threshold

Our previous result shows that PPI interfaces encode information about two broadly distinct topologies, one where the interactor is soluble and another that is fully embedded in the membrane. However, it remains unclear whether the interface features also encode information that can distinguish subtle differences between topologies, such as those of partially embedded interfaces in the membrane (Figure 1c).

To explore this question, we varied the TM coverage threshold *t* from 0.10 to 0.90 in steps of 0.10. For each threshold, PPI interfaces were divided into two topologies. Here, “full-TM” denotes interfaces in which both interacting proteins exceed the threshold *t*, whereas “partial-TM” denotes interfaces in which only one interacting protein exceeds the threshold (Figure 1c, Figure 2b). These labels hence depend on the threshold *t*. For example, at *t* = 0.20, an interface is defined as full-TM if both interacting proteins have TM coverage > 0.20, and as partial-TM if only one interacting protein does and the other is below the threshold *t*. Figure 2b shows AlphaFold predicted example structures for full-TM and partial-TM topologies.

To investigate the topologies with the different thresholds, we apply the same analysis as in the previous subsection. To account for the decreasing sample sizes at different thresholds, we additionally assessed feature importance computed from 50 random subsamples (Methods 4.7).

At a low threshold (*t* = 0.10), the topologies are similar to the comparison between soluble versus fully embedded, and so they were easily distinguished (Acc above 90%, AUC above 0.96). This suggests that the topologies differ strongly in their interface features at this threshold. At a higher threshold, such as t=0.50, the distinction between the topologies was weaker but remained pronounced (72–77% Acc, AUC 0.81–0.84). At the maximum threshold (*t* = 0.90), the two topologies were hard to distinguish (Table 1), which is consistent with the increased overlap of their interface features (Figure 4a). Thus, the greater the difference of the TM coverage between two topologies, the easier is to distinguish between the interfaces (Figure 4b).

**Figure 4.**
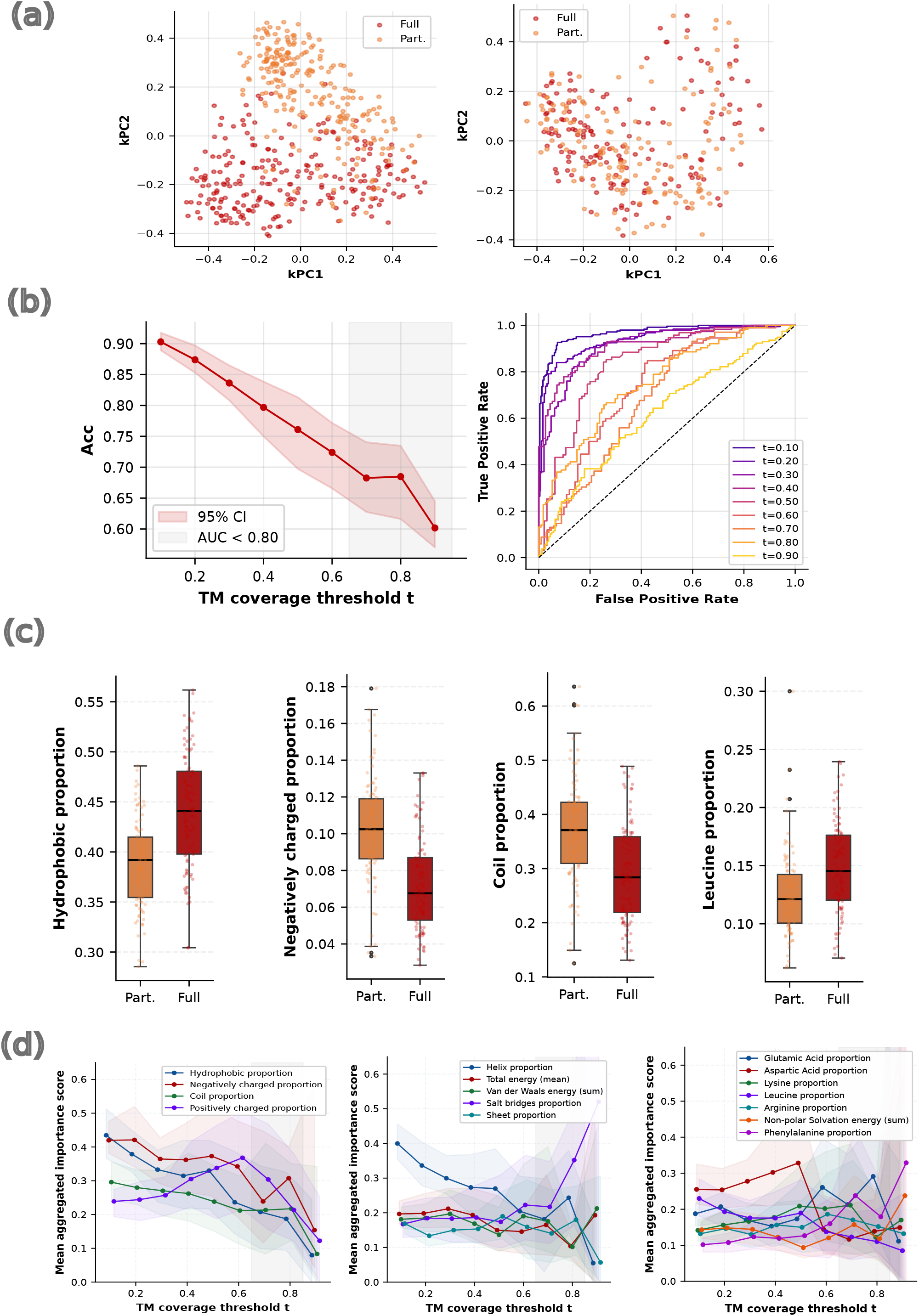
Comparative analysis of partial-TM (Part., orange) and full-TM (Full, red) interfaces across varying TM coverage thresholds. a) Kernel PCA of interface feature space at low (left) (t=0.10) and high (right) (t=0.90) TM coverage thresholds. b) RF Acc across TM coverage thresholds (left, mean *±* 95% CI across 100 subsampling runs per threshold and ROC curves for all thresholds (right) showing AUC (single run). c) Boxplots of the top four most important features comparing partial-TM (Part.) and full-TM (Full) interfaces at t=0.50. d) Mean aggregated feature importance scores across TM coverage thresholds *t* for the most important features (mean *±* 95% CI across 50 random subsampling runs per threshold). For all boxplots in this figure, lower and upper hinges of box plots correspond to the 25th and 75th percentiles, respectively. Lower and upper whiskers extend from the hinge to the smallest or largest value no further than the 1.5*×* interquartile range from the hinge, respectively. Black line represents the median and the black dots represent outliers. All features shown are significant after FDR correction (adjusted p < 0.05).

To gain understanding about the features that characterized differences between these topologies, we identified the interface features that contributed most to the analysis at each value of the parameter *t*. At low values of *t*, the distinction between full-TM and partial-TM topologies is dominated by broad compositional and structural features, particularly the proportion of hydrophobic and negative charge amino acids, as well as coil-coiled and *α*-helix structure elements. At low values of *t*, full-TM interface features have higher proportions of hydrophobic, leucine, and *α*-helix than partial-TM interfaces. In contrast, partial-TM interfaces have higher proportions of negatively charged amino acids and coil-coiled structure elements (Figure 4c, Figure S4).

In contrast, at higher thresholds, partial-TM interfaces were enriched in the proportion of hydrophobic and *α*-helical residues, and had lower proportions of negatively charged and coil-coiled secondary structure elements (Figure S4). Consequently, in distinguishing between topologies, the proportions of hydrophobic, amino acid, and secondary structure elements were less important at higher than at lower values of *t*. Moreover, at high values of *t* (*t* = 0.90), amino acid composition and secondary structure contributed little to distinguish between the topologies, with the exception of the proportion of phenylalanine. The remaining differences were mainly energy related (i.e., total binding energy, van der Waals energy, and non-polar solvation energy) (Figure S4).

Interestingly, at intermediate values of *t*, the proportion of aspartic acid became predominant, whereas the proportions of lysine and arginine remained important up to *t* = 0.60. The proportion of salt bridges was relatively stable and high, at lower and higher values of *t*, respectively (Figure 4d). Overall, this suggests that once the broad differences in membrane embedding become smaller at higher values of *t*, the remaining differences between full-TM and partial-TM interfaces are driven by charged residues and by the proportion of salt bridges.

In summary, these results show that the physicochemical differences in the interface features between full-TM and partial-TM interfaces are smaller at higher TM coverage thresholds. Once both interacting proteins are fully embedded in the membrane at the interface (*t ≥* 0.90), features alone provide limited information to distinguish them based on the coverage threshold. Therefore, we next investigate whether other topological property provide information to distinguish interfaces within this fully membrane-embedded dataset.

### 2.5 Interface features encode the number of TM domains of the interactor

The previous analysis showed that the features of PPI interfaces contain information about subtle topological differences in membrane embedding. In this subsection, we turn to another topological property of TM PPI interfaces, namely the number of TM domains, specifically whether the interactor of the SLC has a single versus multiple TM domains (Figure 1d).

To test this, we focus on a subset of PPIs in which both proteins are fully membrane-embedded (*t* >=0.90). We compare interfaces involving interactors with one TM domain (*n* = 195) with interfaces where the interactor contains more than one TM domain, a set that spans a wide range of TM domains (2–38, median = 6, *n* = 234) (Figure 5a). Example structures for these topologies are shown in Figure 2b. Single-pass and multi-pass interfaces were spread across a coverage range *tleq* 0.90, with most PPI interfaces falling between 0.98 and 1.0 coverage (Figure S5). To explore which features distinguish these topologies, we applied classification, and feature importance analysis (Figure 5).

**Figure 5.**
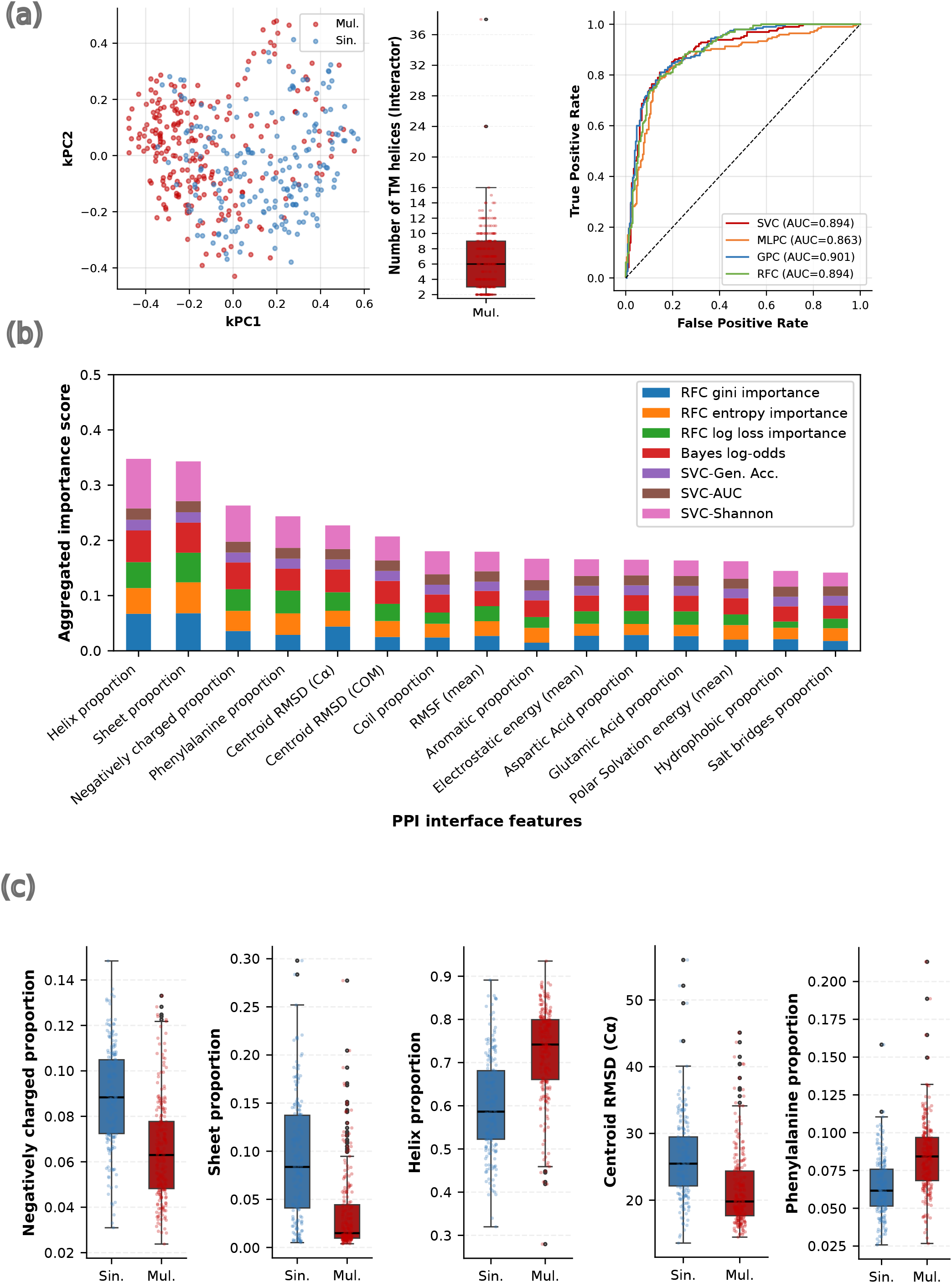
Analysis to distinguish single-pass (Sin., blue) and multi-pass (Mul., red) Interactor. a) Kernel PCA of interface feature space (left). Box-plot (middle) indicates the distribution of TM segments for multi-pass Interactors as reported in UniProtKB. ROC curves for four classifiers (right) reported as AUC. (b) Aggregated feature importance scores for the top 15 interface features across seven importance metrics. c) Box-plots of top-ranked importance features. Blue: Single-pass (Sin.); Red: Multi-pass (Mul.) interfaces (TM). For all boxplots in this figure, lower and upper hinges of box plots correspond to the 25th and 75th percentiles, respectively. Lower and upper whiskers extend from the hinge to the smallest or largest value no further than the 1.5× interquartile range from the hinge, respectively. Black line represents the median and the black dots represent outliers. All features shown are significant after FDR correction (adjusted p < 0.05).

Although both topologies are fully membrane-embedded, they remained clearly distinguishable (Acc 81–83%, AUC 0.87–0.90; Table 1). This indicates that PPI interface features contain substantial information about the interactor TM domain topologies. The most important interface features that characterize these topologies were the proportions of *α*-helix and *β*-sheet, followed by the proportions of negatively charged residues, phenylalanine, and interface geometry. Energy features contributed comparatively little (Figure 5b).

Interfaces involving multi-pass interactors contained higher proportions of *α*-helices and phenylalanine residues. In contrast, interfaces involving single-pass interactors contained more *β*-sheet and coil structure, together with a higher proportion of negatively charged residues (Figure 5c, Figure S6).

The interface geometry also differed between the topologies. Interfaces involving single-pass interactors were more flexible (i.e., higher RMSD, Figure S6). Together, these observations are consistent with the structural organization of PPIs involving multi-pass interactors, which form tightly packed TM *α*-helix bundles.

Together, these observations show that single-pass and multi-pass proteins can be distinguished by information encoded in their physicochemical features. Notably, the local interface alone is sufficient to identify the overall TM architecture of the interactor.

### 2.6 TM interface topologies generalize beyond the SLC interactome

To assess whether the TM interface topologies observed in the SLC interaction dataset are specific to SLC interactions or reflect a broader property of TM protein interactions, we compared them with an independent and large dataset of structurally resolved human PPIs (hereafter referred to as the reference dataset; Methods 4.4). Out of 9, 100 protein complexes filtered from this reference dataset, 2, 831 PPIs contained at least one protein annotated with a TM segment in UniProtKB. Interfaces with a TM coverage *t* > 0.50, on both interacting proteins were the most common topology in both our SLC dataset and the reference dataset, accounting for 58.6% of PPIs (604 of 1, 031) in our SLC dataset and 73.9% (2, 093 of 2, 831) in the reference dataset.

Interfaces with TM coverage *t ≤* 0.50, on both interacting proteins accounted for 32.2% of PPIs in our SLC dataset and 12.4% in the reference dataset. The two asymmetric topologies, in which one interacting protein had TM coverage *t* > 0.50 and the other *t ≤* 0.50, accounted for 1.5% vs. 5.2% and 7.8% vs. 8.5% of PPIs, respectively (Figure 2c, Figure S7).

Together, these distributions show that highly membrane-embedded interfaces (t>0.50) are the predominant topology in both datasets, suggesting that this is a general property of TM protein interactions rather than a feature specific to the SLC interactome. One possible explanation is that interfaces with higher TM coverage are energetically favored. Indeed, higher TM coverage was associated with greater hydrophobicity and more favorable interaction energies in the SLC dataset.

## 3 DISCUSSION

In this work, we asked whether physicochemical features of the interfaces formed by transmembrane protein-protein interactions (PPIs) contain information about the interface topology, which we defined as the interface position and arrangement with respect to the membrane. We studied a large and representative dataset of PPIs derived from the human SLC interactome. We found that interface features can distinguish between soluble interfaces versus those fully embedded in the membrane. Similarly, interface features were able to capture substantial information to distinguish between the topology of interfaces with different degrees of membrane embedding, as well as interfaces involving interacting proteins with single versus multiple TM domains. Together, these findings demonstrate that the features of PPI interfaces retain a substantial amount of information about the physicochemical constraints imposed by the environment in which they form.

The clearest differences between features were observed between soluble interfaces and those fully embedded in the membrane. The distinction became progressively weaker as the interacting proteins became more similar in their degree of membrane embedding. The observed changes in amino acid composition and secondary structure are consistent with the energetic constraints imposed by the lipid bilayer (von Heijne 1992, Cymer et al. 2015). The same interface features remained informative when the analysis was restricted to interactors with at least one TM domain, indicating that these differences reflect the local membrane environment at the interaction interface rather than simply the presence of a TM domain elsewhere in the protein.

In addition to membrane embedding, our results indicate that interface features also contain information about TM architecture, here defined as the number of TM domains per protein. Interface features can distinguish single-pass from multi-pass interactors, consistent with expected differences in TM-domain packing and conformational flexibility between proteins of single versus multi-pass (Walters and DeGrado 2006). Multi-pass interfaces show more aromatic residues, in line with the known role of aromatic and aliphatic residues in packing TM *α*-helices (Mayol et al. 2019). Overall, these observations suggest that interface features capture not only the extent of membrane embedding but also structural differences between TM architectures.

The similar distributions of membrane embedding observed in a reference dataset suggest that the interface topologies studied in the interactome of solute carrier transporters is representative of other larger TM PPIs datasets, and most likely, unbiased. More generally, our results indicate that different topologies and TM architectures leave a measurable signal on PPI interfaces that can be captured by physicochemical interface features. This information may support studies of membrane protein interaction topology and could improve the interface annotation of predicted protein complexes, as well as guide protein design strategies focusing on TM PPI. It was shown before, that AlphaFold can be used to identify interaction interfaces, using either the full length protein or protein fragments, however it remains challenging to experimentally validate the interaction interfaces (Burke et al. 2023, Lee et al. 2024).

Future work should evaluate these observations in larger and more diverse datasets as additional membrane protein complexes are experimentally resolved and thousands are predicted with AlphaFold and other structural prediction tools. Moreover, molecular dynamics simulations in explicit lipid bilayers will further help to investigate the interplay between membrane biophysics and the topology of protein-protein interaction interfaces.

## 4 METHODS

### 4.1 Structure Prediction and Molecular Dynamics Simulations

Protein interactions were obtained from the SLC transporter interactome dataset (Frommelt et al. 2025). Complex structures were predicted based on AlphaFold v3.0 (AF3) Abramson et al. (2024) with one seed per interaction, generating five ranked models per PPI. Models were filtered using confidence thresholds of ipTM *≥* 0.3 and predicted DockQ (pDockQ) *≥* 0.23 (Zhu et al. (2023)). Low-confidence terminal regions were removed by iterative trimming of N- and C-termini until the mean pLDDT exceeded 70 across a five-residue sliding window (Schweke et al. (2024)). We generated strucutral models for all 19,000 experimentally obtained PPIs using AF3. After filtering we came up with 2,055 structural models for further analysis.

Simulations were performed using AMBER2024 (Case et al. (2023)) with pmemd.cuda (Salomon-Ferrer et al. (2013)) and the ff14SB force field (Maier et al. (2015)). Each complex was prepared with pdb4amber and ParmEd, solvated in a truncated octahedral TIP3P water box with neutralizing Na^+^ and Cl^−^ ions under periodic boundary conditions, PME electrostatics, and SHAKE hydrogen constraints (Ryckaert et al. (1977)). Each system underwent energy minimization (40,000 steps), restrained NVT heating from 0 to 300 K (Langevin thermostat, 5 ps^−1^, 1 fs timestep), NVT equilibration (500 ps), and NPT equilibration (500 ps) with gradually released restraints at 2 fs, followed by a 10 ns NPT production run at 300 K and 1 bar (Langevin thermostat, 2 ps^−1^; Berendsen barostat) with a 4 fs timestep enabled by HMR, saving 100 frames per complex. Binding free energies were estimated by MM-GBSA on all 100 frames (Genheden and Ryde (2015)). Residue-level features were extracted using BioPython (Cock et al. (2009)), CPPTRAJ (Roe and Cheatham (2013)), and DSSP (Kabsch and Sander (1983)). To determine contact residues, GetContacts (Venkatakrishnan et al. (2019)) was used, to get contact information for all non-covalent interaction types.

### 4.2 Feature Engineering and Aggregation

Interface residues were defined using a 10 Å residue distance cutoff (Collins et al. 2024). Only contacts observed in more than 30% of MD frames were retained as stable, excluding transient interactions (Jaeger-Honz et al. 2024).

The interface-level features were aggregated per PPI and grouped into six categories: amino acid composition, secondary structure, energy, geometry, flexibility, and intermolecular contacts. Interface length was defined as the total number of residues of both proteins at the interaction interface.

**Amino acid composition** features captured residue-level chemical properties using proportions of all 20 amino acids and six broader physicochemical classes. Each category was computed as smoothed relative frequencies within the interface. Additive smoothing was applied per category using a fixed pseudocount (*α* = 1.5), with a corresponding normalization offset applied to maintain consistent scale of regularization across feature groups. This smoothing prevents zero-valued features and ensures numerical stability under subsequent logit transformation (Supplementary Methods).

**Secondary structure** features quantified interface structural organization. DSSP states were grouped into three classes: *α*-helix (H, G, I), sheet (E), and coil (B, T, S, C), with the same smoothed relative frequency encoding applied.

**Energy** features summarized MM/GBSA van der Waals, electrostatic, polar solvation, and nonpolar solvation terms (Genheden and Ryde 2015), aggregated as both the mean and sum across interface residues to capture per-residue intensity and cumulative interface-level contribution, respectively. Total binding free energy and its standard deviation represented overall binding strength and energy variability.

**Geometric** features quantified interface shape and spatial organization. Compactness was measured as the root-mean square distance of interface residues from their geometric centroid (Kufareva and Abagyan 2012). Planarity was measured as the root-mean square distance to a best-fit plane obtained by singular value decomposition (SVD) (Rodrigues et al. 2022, Jones and Thornton 1996). Both were calculated using C*α* atoms and residue centers of mass (COM) (Supplementary Methods).

**Flexibility** was quantified using the mean and summed root-mean square fluctuation (RMSF) of interface residues.

**Contact-based** features summarized stable intermolecular interactions across seven classes: hydrogen bonds, salt bridges, *π*-stacking, *π*-cation interactions, T-stacking, van der Waals contacts, and water bridges. Each class contributed a raw count and an interface-length-normalized rate, computed as the smoothed contact count divided by total interface length using the same pseudocount regularization scheme. Additional features included total contact count, contact density, interface length, and interface fraction (Supplementary Methods). Details on amino acid grouping, secondary structure definitions, and raw GetContacts interaction-type mappings are provided in Supplementary Tables S2-S4.

Before classification, count-based features were log(*x* + 1)-transformed to preserve zero counts and reduce skew. Geometric features and MM/GBSA standard deviations were log-transformed (Box and Cox 1964) to reduce skewed distributions. Proportion-based features were logit-transformed to map bounded values onto an unbounded scale (Warton and Hui 2011). Binding free energy mean and sum features, as well as RMSF-based flexibility measures, were retained on their original scale because they preserve physically meaningful signed or magnitude-based values, and additional transformation would reduce interpretability without improving numerical stability. All features were standardized to zero mean and unit variance prior to classification (Pedregosa et al. 2011).

### 4.3 Transmembrane Domain Annotation

To classify our data by topology and support further analysis, we introduced a TM coverage score per protein of each PPI. Therefore, TM domain annotations were retrieved from UniProtKB (UniProt Consortium 2023). In UniProtKB, these annotations are based on experimental evidence, homology, or computational predictions (TMHMM) (UniProt Consortium 2017).

For each protein in a PPI, TM coverage was computed as the maximum proportion of residues within an annotated TM segment that participate in the PPI interface, across all segments. We used the maximum coverage value to represent the segment most involved in the interface. This yielded per-protein coverage scores for the SLCs and the interactors for each distinct PPI.

In addition, the number of annotated TM domains per interactor was recorded. Interactors with a single TM domain were defined as singlepass, whereas those with two or more TM domains were defined as multi-pass.

### 4.4 Reference dataset

To assess whether the observed TM coverage distributions are specific to the dataset, which is used in this study we used a structurally resolved human PPI dataset (Burke et al. 2023). Structures were filtered to pDockQ > 0.23 scores, yielding 9,100 unique structures. PPI AF3 interfaces were defined using the same 10 Å distance cutoff between residues of the interacting proteins, as used in the SLC dataset (Frommelt et al. 2025) (Section 4.2). TM interface residues were mapped using the same UniProtKB-based annotation procedure (Section 4.3).

### 4.5 Supervised Machine Learning

To visualize class separation in interface feature space, we applied kernel principal component analysis (PCA) with a radial basis function (RBF) kernel to the standardized features, projecting each interface onto its top two components (Schölkopf et al. 1998). To evaluate the predictive value of aggregated features, we employed four complementary classifiers from scikit-learn (Pedregosa et al. (2011)) to capture a range of decision boundary geometries and uncertainty estimates. Support Vector Classifier (SVC) (Vapnik (1998)), Multilayer Perceptrons (MLP) (Hornik (1991)), Random Forests (RF) (Breiman (2001)), and Gaussian Process Classifiers (GPC) (Rasmussen and Williams (2006)), to capture a range of decision boundary geometries. Performance was assessed using 5*×*5 nested cross-validation to ensure that predictions are unbiased. The outer loop estimates generalization on held-out data while the inner loop performs hyperparameter optimization within training folds, preventing information leakage (Varma and Simon 2006).

### 4.6 Performance Metrics

Supervised classification assessed whether the label is encoded in the extracted interface features.. Performance is evaluated using Accuracy (Acc), which measures the fraction of correctly predicted labels (Hand et al. (2024)). To assess performance, we used Receiver Operating Characteristic (ROC) curves by calculating sensitivity and 1-specificity for each threshold. The Area Under the Curve (AUC) quantifies overall discriminative performance (Hanley and McNeil (1982)). To quantify reliable information content in predicted posterior probabilities we computed a modified mutual information (MI)(MacKay (2003)), Wöber et al. (2021)). Contributions of misclassified samples in the MI are negated to penalize wrong predictions. Statistical significance against a majority-class baseline was assessed using the McNemar’s test (McNemar (1947)). For RF classifiers in the downsampled comparisons, performance was additionally assessed across 100 independent random balanced subsampling runs (without replacement), with Acc and AUC reported as mean *±* standard deviation (Kohavi 1995).

### 4.7 mFeature Information Analysis

To quantify how individual features contribute to classification, the aggregated features were evaluated with seven complementary metrics.

RF impurity-based importance using Gini, entropy, and log loss criteria was extracted from a model trained on the full feature set (Breiman 2001, Louppe 2013). These values sum to one by definition and reflect each feature’s contribution to impurity reduction across the ensemble. For Bayesian evaluation, per-feature Gaussian Process Classifiers were compared against a constant baseline using Bayes Factors in log-odds form, where larger values indicate stronger evidence for feature inclusion (Rasmussen and Williams 2006, Kass and Raftery 1995). Discriminative power was assessed by training per feature SVCs and evaluating Acc, AUC (Hanley and McNeil 1982), and modified MI (MacKay 2003). Each metric was then divided by its sum across all features to yield relative proportions that are comparable between methods. Features were finally ranked by their sum across all seven criteria, where higher values reflect more consistent importance.

For the comparison between interfaces with high and intermediate TM coverage, feature importance was assessed across 50 independent random balanced subsampling runs per threshold (t=0.10–0.90), each drawn without replacement, with mean values and 95% confidence intervals (CI; 2.5th and 97.5th percentiles) reported across runs (Kohavi 1995).

Mann-Whitney U test p-values for feature comparisons were corrected for multiple testing using the Benjamini-Hochberg (FDR) procedure (Benjamini and Hochberg 1995).

## 7 DATA AVAILABILITY STATEMENT

The aggregated feature matrix and code underlying this article are available on GitHub at https://github.com/lallm/SLC-topologies. The raw feature data before aggregation and the reference dataset are available on Zenodo at https://doi.org/10.5281/zenodo.21306901.

## AUTHOR CONTRIBUTIONS

**Lisa Allmesberger-Riegler:** Conceptualization; Data curation; Formal analysis; Investigation; Methodology; Software; Validation; Visualization; Writing – Original Draft Preparation; **Fabian Frommelt:** Conceptualization; Data curation; Formal analysis; Funding Acquisition; Investigation; Software; Visualization; Writing – Original Draft Preparation; **Brianda L. Santini:** Data curation; Formal analysis; Investigation; Software; **Giulio Superti-Furga:** Funding acquisition; Project Administration; Resources; Supervision; **Evandro Ferrada:** Funding Acquisition; Investigation; Methodology; Software; Supervision; Validation; Writing – Original Draft Preparation; **Peter Sykacek:** Funding Acquisition; Investigation; Methodology; Project Administration; Resources; Supervision; Writing – Original Draft Preparation

## 9 ACKNOWLEDGMENTS

This research was funded by the Vienna Science and Technology Fund (WWTF) and the City of Vienna through grant 10.47379/LS23028 mlDIAMANT. The authors thank the CeMM IT team, especially Patricia Carey, for their support during the development of the structural prediction pipeline.

## 10 AI DISCLOSURE STATEMENT

During the preparation of this manuscript, the authors used in part AI-assisted technologies to support code development and data analysis scripts. After using these tools, the authors reviewed, edited, and take full responsibility for the final content and technical accuracy of the manuscript.

## 11 SUPPLEMENTARY MATERIAL DESCRIPTION

### Supplementary Figures

- **Figure S1**. Additional box plots for soluble vs. TM interface features.
- **Figure S2**. Results of TM vs. TM not at the interface (t=0).
- **Figure S3**. Box plots differentiating fully embedded vs. not embedded (t=0) topologies.
- **Figure S4**. Mean values of interface features across TM coverage thresholds.
- **Figure S5**. Distribution of single- and multi-pass topologies among fully embedded PPIs.
- **Figure S6**. Box plots differentiating single-pass vs. multi-pass topologies.
- **Figure S7**. Scatter plot of the TM coverage at the PPI interface for the reference data set (Burke et al. 2023).

### Supplementary Tables

- **Table S1**. Overview of feature categories, definitions and applied transformations.
- **Table S2**. Amino Acid classes definition.
- **Table S3**. Secondary Structure grouping.
- **Table S4**. Contact types from GetContacts and specific grouping.

### Supplementary Methods

- **Feature Aggregation** Additional description of employed feature aggregation procedures.

